# M1 recruitment during interleaved practice is important for encoding, not just consolidation, of novel skill memory

**DOI:** 10.1101/2023.07.21.550118

**Authors:** Taewon Kim, Hakjoo Kim, Benjamin A. Philip, David L. Wright

## Abstract

Primary motor cortex (M1) plays a major role in motor memory acquisition and retention in humans, but its role in interleaved practice (as opposed to repetitive practice) remains unknown. We anticipated that the improved retention typically associated with interleaved practice depends on M1, and thus cathodal transcranial direct current (ctDCS) stimulation to M1 during training would disrupt this improved retention. The benefits of interleaved practice have been reported to occur from more effective consolidation, manifested as rapid skill memory stabilization followed by more long-term enhancement. While we observed the expected decline in retention performance following interleaved practice paired with ctDCS, this reduced retention resulted from more modest encoding of novel skill memory during acquisition rather than from disruption of offline consolidation processes. These data highlight the broad role played by motor cortex for both encoding and retention of novel skill memory.

## MAIN CONTENT

Learning motor skills is central to everyday human functioning as well as a common feature of motor rehabilitation ^1,2^. The neural mechanisms of motor learning have been well studied and depend in part on both online acquisition and offline consolidation processes involving the primary motor cortex (M1), which occur during periods of wake ^3-5^ or sleep ^6,7^ respectively. Changes in neural excitability in M1 occur throughout the acquisition of novel skills in the intact brain as well as during the restoration of motor function in the damaged brain ^8,9^. Thus, understanding neuroplastic change at M1 is critical for understanding skill learning as well as potentially offering insights into novel therapeutic approaches for motor recovery.

It is generally accepted that increased practice is crucial for motor learning. However, the effect of practice depends on more than just its amount: skill retention also depends on practice schedule^10^. For example, repetitive practice (RP) is a schedule that involves extensive repetition of a novel skill before exposure to the acquisition of subsequent skills; RP invokes a moderate level of challenge and fosters rapid performance gains but hinders retention. In contrast, interleaved practice (IP) is a relatively more demanding practice format because it consists of frequent switching between to-be-learned skills during training; IP is associated with a slow rate of skill acquisition but leads to improved long-term retention^11,12^. Improved skill memory following IP has been ascribed to more effective offline consolidation, which is associated with elevated post-practice M1 excitability ^5^. Moreover, neuroimaging data reveals that IP is associated with the emergence of more extensive task-related functional connectivity of sensorimotor and frontoparietal networks when compared with RP ^13^. A causal role for M1 for enhanced novel skill retention is highlighted by the finding that administration of anodal (excitatory) transcranial direct current stimulation (atDCS) at M1 during RP promotes time and sleep-associated consolidation typically observed after IP ^14-16^. However, these findings have only demonstrated that M1 can play a contributory role in skill retention but do not demonstrate that M1 plays a necessary role.

In the present investigation, we directly tested M1’s causal role by down-regulating it during IP via cathodal (inhibitory) transcranial direct current stimulation (ctDCS) to determine if the performance benefits of IP could be disrupted. Based on previous work, we anticipated that, compared to the administration of sham tDCS, reducing the excitability of M1 during IP should impede post-practice consolidation, thus reducing the offline benefit traditionally observed after IP ^5,16^.

Fifty-four right-handed undergraduate students (ages 19-23) were randomly assigned to one of three groups (N = 18/group): RP-Sham, IP-Sham, and IP-ctDCS. All groups practiced three unique 6-item, motor sequence learning task (specifically, a discrete sequence production task) with their left index finger ^16-18^. All participants experienced nine blocks of 21 trials resulting in 189 total trials of practice. For RP, each block of practice only included practice with a single task. In contrast, participants assigned to IP practice all three tasks in each block. Training was preceded by a baseline test and followed by three retention tests at 5 min, 6 h, and 24 h after the completion of practice (see Fig. 1). All test blocks included 21 trials: 7 trials of each task presented in an RP format. The primary outcome measure was total response time (TT), with lower TT representing better performance.

**Fig. 1:**
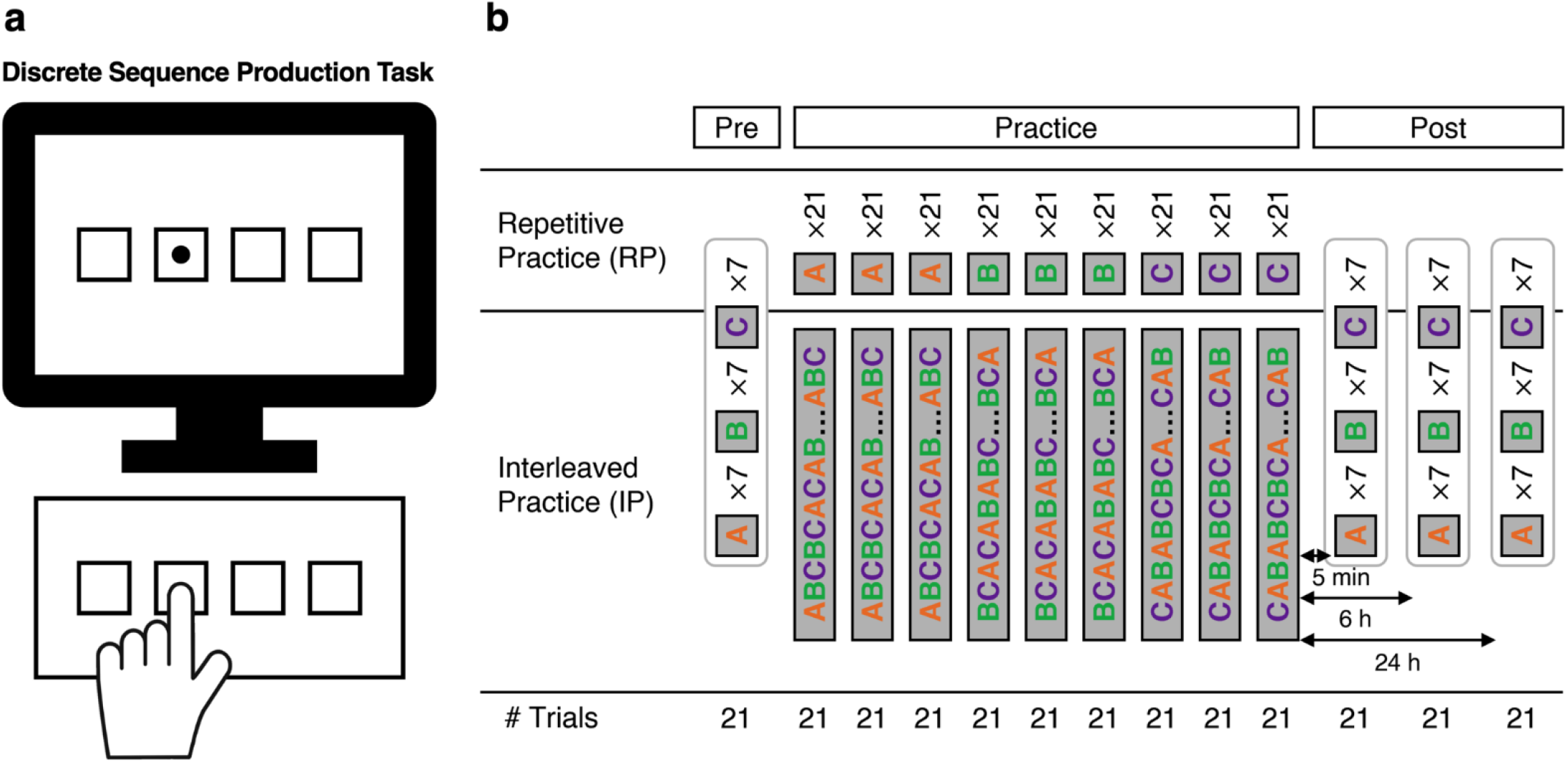
The experimental procedure. **(a)** Motor sequence learning task (specifically, a discrete sequence production task) was used ^16-18^, each of which required the execution of six key presses with the left index finger only that was directed by the presentation of a visual signal on the computer display. **(b)** During the practice phase, individuals in the IP-ctDCS group were administered cathodal stimulation at the right M1. Practice included nine training blocks of 21 trials organized in either a repetitive or an interleaved format. Tests were administered prior to training (Pre) and following training at Post_5min_, Post_6h_, and Post_24h_. All tests consisted of 21 trials that included seven trials of each of the three sequences in a repetitive format. Stimulation (real or sham) was only present during the training blocks.

We confirmed IP’s previously identified benefits (vs. RP) for offline gain^16,17,19^ via a 2 (Group: IP-Sham, RP-Sham) × 3 (Time Point: Post_5min_, Post_6h_, Post_24h_) analysis of variance (ANOVA) with repeated measures on the last factor. We found a significant interaction effect, *F*(2, 68) = 8.39, *p* < .001, η_p_^2^ = 0.2. While TT was similar 5 min after training (t(34) = 1.29, *p* = .2, *d* = 0.29), TT was significantly lower at Post_6h_ (t(34) = 3.81, *p* < .0001, *d* = 0.78) and Post_24h_ (t(34) = 7.06, *p* < .0001, *d* = 1.4) for the IP-Sham compared to the RP-Sham group (see Fig. 2a). Congruent with previous work ^16^, IP-Sham participants exhibited stable performance across the initial 6 h after training (t(17) = 0.45, *p* = .62, *d* = 0.11) and a reduction in TT following overnight sleep (t(17) = 3.74, *p* < .0001, *d* = 0.91). In contrast, the RP-Sham group displayed significant forgetting in the first 6 h after training (t(17) = 2.98, *p* < .01, *d* = 0.72) and no offline effects during the subsequent 18 h that included overnight sleep (t(17) = 0.94, *p* = .349, *d* = 0.23). These and other effects could not be explained by differences in baseline performance because the three groups performed similarly at baseline, *F*(2, 51) = 0.32, *p* = .73, η_p_^2^ = 0.01.

**Fig. 2:**
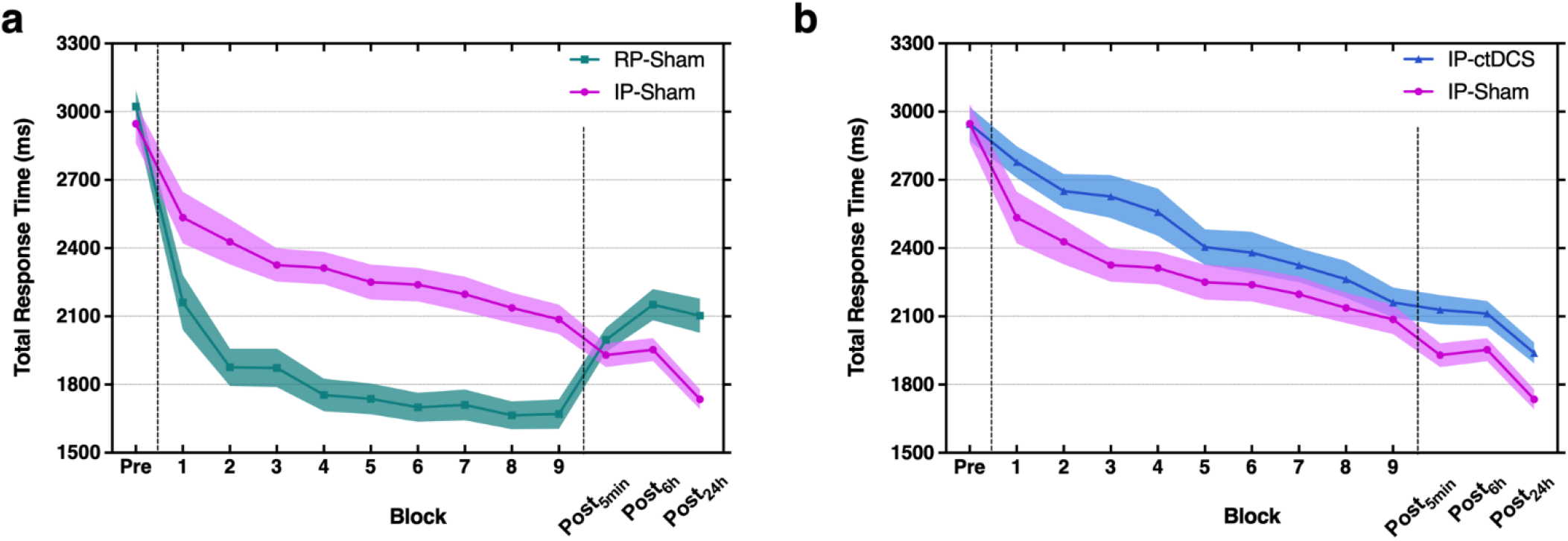
Total response time (ms) for baseline (Pre), training, and post-practice time points (Post_5min_, Post_6h_, Post_24h_). **(a)** IP led to greater long-term performance due to offline improvement in the immediate (5 min), medium-term (6 h), and long-term (24 h) compared to the final block of practice, whereas RP shows significant forgetting at all post-practice recall time points compared to the end of practice. This replicates previous findings on the influence of practice structure on learning novel motor skills ^11,14,16,17,19^. **(b)** IP-ctDCS at M1 disrupted the encoding of a motor sequence memory for later offline consolidation. This was reflected in IP-ctDCS group showing significantly greater TT than IP-Sham for all training blocks except the final one. However, inhibitory ctDCS to M1 did not prevent subsequent offline consolidation, as shown by continuing post-training decreases in TT (i.e., performance improvements) for both IP groups in the medium-long term (6-24 h).

We identified the effects of ctDCS during IP by 2 (Group: IP-Sham, IP-ctDCS) × 9 (Block) ANOVA. We found a significant interaction, *F* (8, 272) = 2.26, *p* < .05, η_p_^2^ = 0.06), resulting from higher TT for the IP-ctDCS participants compared to their IP-sham counterparts for all training blocks except for Block 9. The impact of poorer encoding as a result of ctDCS during IP training remained throughout retention, reflected in a significant main effect of Group, *F*(1, 68) = 7.58, *p* < .01, η_p_^2^ = 0.1, for the test blocks (*p* < .001 for all test blocks) revealing poorer novel skill memory for the IP-ctDCS group. While supplementing IP-ctDCS disrupted memory development compared to IP-sham, ctDCS had no impact on subsequent consolidation, as indicated by the significant main effect of Test Block, *F*(2, 68) = 35.51, *p* < .001, η_p_^2^ = 0.51. Specifically, TT remained stable across the first 6-h following training (t(35) = 0.13, *p* = .9, *d* = 0.02) but was significantly lowered at the 24-h test following sleep (t(35) = 7.23, *p* < .0001, *d* = 1.2) (see Fig. 2b). The absence of a Group × Test Block interaction, *F* (2, 68) = 0.44, *p* =0.65, η_p_^2^ = 0.01) verified that the nature of consolidation commonly observed following IP occurred irrespective of the presence or absence of ctDCS during training.

Our results reveal that M1 has a broader influence on memory development and retention than previously recognized^5,14,16^. Here, we demonstrate that direct downregulation of M1 via cathodal stimulation during IP impedes the construction of a novel motor memory yet preserves effective consolidation. This complements earlier work that showed enhanced consolidation but unchanged acquisition when supplementing RP with anodal tDCS to M1^16^. Together, these results highlight that a complex interplay between the physical practice format and the type of non-invasive brain stimulation determines the observed online and offline behavioral outcomes, with M1 involved in multiple mechanisms. The contribution of M1 to skill memory encoding is similar to that assigned to the dorsal premotor cortex (PMd) in the context of RP in previous work ^17^. These data are congruent with proposals that both M1 and PMd are important for development of motor sequence representation across training ^20,21^.

## METHODS

### Participants

Participants assigned to IP with cathodal stimulation at the right M1, and two sham conditions, each including a separate group of 18 participants, received training in either an RP or IP format. Real (IP-ctDCS) or sham (RP-Sham, IP-Sham) stimulation was applied during the entire 20-min period of practice. Participants were blinded to the stimulation condition. All participants completed an informed written consent approved by Texas A&M University’s Institutional Review Board before any involvement in the experiment.

### Transcranial direct current stimulation (tDCS)

We targeted the right M1 with a 1 × 1 tDCS electrode montage. Real stimulation consisted of a 2 mA current applied via a 25 cm^2^ (5 × 5 cm) cathode and a 35 cm^2^ (5 × 7 cm) reference electrode covered by saline-soaked sponges, resulting in a maximum current density of 0.08 mA/cm^2^ administered using a 9 V battery-driven stimulator (tDCS Stimulator; TCT Research Limited, Hong Kong). The cathode was located at the right M1 (i.e., C4, International 10-20 system) and was paired with a reference electrode above the left supraorbital region. This placement system has established accuracy for targeting M1, with the current flow simulation shown in Fig. 3. The current flow associated with this electrode montage was modeled using HD-Explore™ (Soterix Medical Inc., New York, NY) and revealed heightened current flow at regions described as right M1 in the human motor area template ^22^. Participants in sham stimulation conditions (RP-Sham, IP-Sham) involved the same electrode configuration, but stimulation was only delivered for 30-sec at the beginning and end of the training period.

**Fig. 3:**
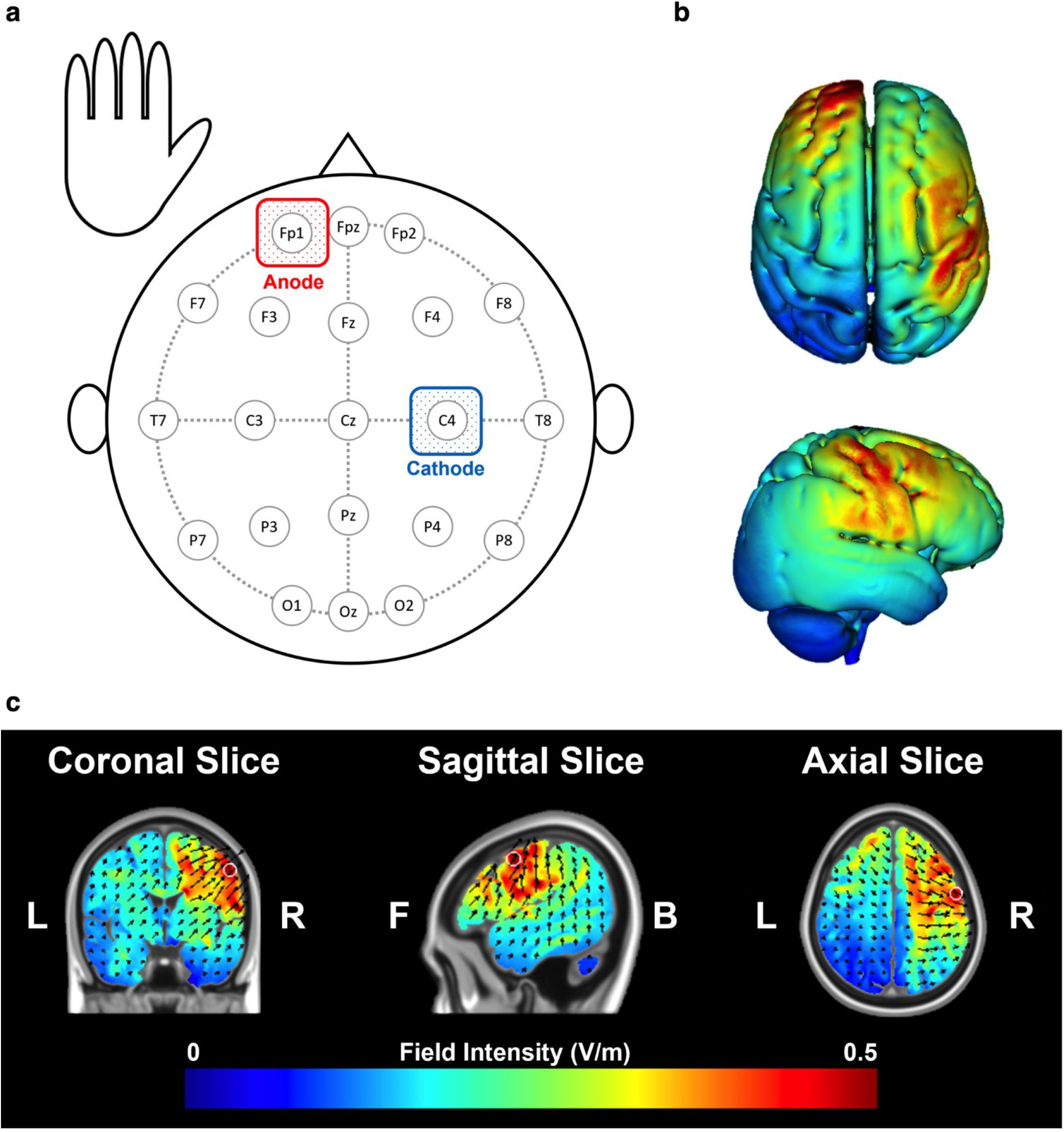
tDCS montage used during practice in IP. **(a)** Cathodal stimulation for the interleaved practice, real tDCS condition involved placement of the cathode 20% of the auricular measurement from Cz (determined based on the International 10–20 system) which placed this electrode above C4 with a reference electrode at the left supraorbital region. Participants in sham stimulation conditions (Repetitive-Sham, Interleaved-Sham) involved the same electrode configuration, but stimulation was only delivered for 30-sec at the beginning and end of the training period. **(b, c)** The current flow associated with this electrode montage was modeled using HD-Explore™ (Soterix Medical Inc., New York, NY) (figure illustrates right M1). Heightened current flow was observed (0.507 V/m) at MNI coordinates (x: 52, y: −1, z: 45). These locations (noted with a white circle) fell within the boundaries described as right-M1, respectively, in the human motor area template ^22^.

### Motor sequence learning task

We employed a motor sequence learning task, specifically, a discrete sequence production task ^18^. To ensure that we studied cortical activity related to motor execution rather than stimulus-response mappings and probabilities ^23,24^, we did not use the four-finger (one per button) design of the traditional serial reaction time task paradigm ^7^. Instead, our participant used only their left index finger for all keys ^16,17,19^, which creates real, non-isometric motor demands in the context of a key pressing task ^25^ (Fig. 1a). A standard keyboard was used. Individuals executed a keypress to a visual signal that was spatially compatible with the position of the key. Once a correct key was pressed, the next visual signal in the sequence was presented. The primary dependent variable assessing motor sequence performance was total response time (TT), which was the interval from the presentation of the first stimulus to the correct execution of the final keypress of the motor sequence.

All individuals completed nine blocks of 21 trials each, totaling 189 trials of practice. These 189 trials were divided between three 6-key motor sequences (63 trials per sequence). The content of each block depended on practice schedule (Fig. 1b). For RP, each block contained 21 repetitions of a single sequence; for IP, each block contained 7 trials of each of the three sequences, presented in counterbalanced order. Test blocks were presented in RP format without stimulation. All participants completed 3 Test blocks at 4 occasions: prior to training (Pre), immediately after training (5 min), 6 h later, and 24 h later (Fig. 1b).

### Statistical analyses

To ensure the normality of the data used in this study, we assessed it using the Shapiro-Wilk test. The test statistic (W) was 0.97, with a corresponding *p*-value of .21. As the *p*-value was greater than .05, we failed to reject the null hypothesis that the data were normally distributed; therefore, we used parametric ANOVA for all analyses. In addition, a Bonferroni adjustment was made when conducting post-hoc multiple comparisons. An initial 3 (Group: RP-Sham, IP-Sham, IP-ctDCS) between-subject ANOVA was conducted to assess if individuals assigned to each of the experimental conditions exhibited similar performance during the baseline test block. The evaluation of online performance (during training) was assessed using two separate two-way ANOVAs with repeated measures on the last factor: (1) Group (IP-ctDCS, IP-Sham) × Block and (2) Group (IP-Sham, RP-Sham) × Block. To determine the effect of ctDCS in M1 and practice structure in offline performance gains, two separate two-way ANOVAs with repeated measures on the last factor were performed: (1) Group (IP-ctDCS, IP-Sham) × Time Point (Post_5min_, Post_6h_, Post_24h_) and (2) Group (IP-Sham, RP-Sham) × Time Point (Post_5min_, Post_6h_, Post_24h_). If we find significant effects, we used Tukey’s Honest Significant Differences post-hoc assessment to identify differences in means.

## References

1 Kitago, T. & Krakauer, J. W. Motor learning principles for neurorehabilitation. Handbook of clinical neurology 110, 93–103 (2013).

2 Krakauer, J. W. Motor learning: its relevance to stroke recovery and neurorehabilitation. Current opinion in neurology 19, 84–90 (2006).

3 Hadipour-Niktarash, A., Lee, C. K., Desmond, J. E. & Shadmehr, R. Impairment of retention but not acquisition of a visuomotor skill through time-dependent disruption of primary motor cortex. Journal of Neuroscience 27, 13413–13419 (2007).

4 Muellbacher, W. et al. Early consolidation in human primary motor cortex. Nature 415, 640–644 (2002).

5 Tunovic, S., Press, D. Z. & Robertson, E. M. A physiological signal that prevents motor skill improvements during consolidation. Journal of neuroscience 34, 5302–5310 (2014).

6 Robertson, E. M., Pascual-Leone, A. & Press, D. Z. Awareness modifies the skill-learning benefits of sleep. Current biology 14, 208–212 (2004).

7 Walker, M. P., Brakefield, T., Allan Hobson, J. & Stickgold, R. Dissociable stages of human memory consolidation and reconsolidation. Nature 425, 616–620 (2003).

8 Dimyan, M. A. & Cohen, L. G. Neuroplasticity in the context of motor rehabilitation after stroke. Nature Reviews Neurology 7, 76–85 (2011).

9 Winstein, C. J. & Kay, D. B. Translating the science into practice: shaping rehabilitation practice to enhance recovery after brain damage. Progress in brain research 218, 331–360 (2015).

10 Guadagnoli, M. A. & Lee, T. D. Challenge point: a framework for conceptualizing the effects of various practice conditions in motor learning. Journal of motor behavior 36, 212–224 (2004).

11 Kim, T., Chen, J., Verwey, W. & Wright, D. Improving novel motor learning through prior high contextual interference training. Acta psychologica 182, 55–64 (2018).

12 Kim, T., Rhee, J. & Wright, D. L. Allowing time to consolidate knowledge gained through random practice facilitates later novel motor sequence acquisition. Acta Psychologica 163, 153–166 (2016).

13 Lin, C.-H. J. et al. Contextual interference enhances motor learning through increased resting brain connectivity during memory consolidation. NeuroImage 181, 1–15 (2018).

14 Wright, D. et al. Consolidating behavioral and neurophysiologic findings to explain the influence of contextual interference during motor sequence learning. Psychonomic bulletin & review 23, 1–21 (2016).

15 Wright, D. L. & Kim, T. Contextual interference: New findings, insights, and implications for skill acquisition. Skill acquisition in sport, 99–118 (2019).

16 Kim, T., Kim, H. & Wright, D. L. Improving consolidation by applying anodal transcranial direct current stimulation at primary motor cortex during repetitive practice. Neurobiology of Learning and Memory 178, 107365 (2021).

17 Kim, T., Buchanan, J. J., Bernard, J. A. & Wright, D. L. Improving online and offline gain from repetitive practice using anodal tDCS at dorsal premotor cortex. npj Science of Learning 6, 31 (2021).

18 Abrahamse, E. L., Ruitenberg, M. F., De Kleine, E. & Verwey, W. B. Control of automated behavior: insights from the discrete sequence production task. Frontiers in human neuroscience 7, 82 (2013).

19 Kim, T. & Wright, D. L. Transcranial direct current stimulation of supplementary motor region impacts the effectiveness of interleaved and repetitive practice schedules for retention of motor skills. Neuroscience 435, 58–72 (2020).

20 Wiestler, T., Waters-Metenier, S. & Diedrichsen, J. Effector-independent motor sequence representations exist in extrinsic and intrinsic reference frames. Journal of Neuroscience 34, 5054–5064 (2014).

21 Wiestler, T. & Diedrichsen, J. Skill learning strengthens cortical representations of motor sequences. Elife 2, e00801 (2013).

22 Mayka, M. A., Corcos, D. M., Leurgans, S. E. & Vaillancourt, D. E. Three-dimensional locations and boundaries of motor and premotor cortices as defined by functional brain imaging: a meta-analysis. Neuroimage 31, 1453–1474 (2006).

23 Lungu, O., Wächter, T., Liu, T., Willingham, D. & Ashe, J. Probability detection mechanisms and motor learning. Experimental Brain Research 159, 135–150 (2004).

24 Philip, B. A., Wu, Y., Donoghue, J. P. & Sanes, J. N. Performance differences in visually and internally guided continuous manual tracking movements. Experimental brain research 190, 475–491 (2008).

25 Chettouf, S., Rueda-Delgado, L. M., de Vries, R., Ritter, P. & Daffertshofer, A. Are unimanual movements bilateral? Neuroscience & Biobehavioral Reviews 113, 39–50 (2020).

